# Predicting human microbe-drug associations via graph attention network with multiple kernel fusion

**DOI:** 10.1101/2023.08.15.553383

**Authors:** Sairu Shi, Shu Kong, Qingwei Zhang, Ji Zhang

## Abstract

Microbial dysregulation may lead to the occurrence of diseases, and using microbe and drug data to infer the microbe-drug association has attracted extensive attention. There have been many studies to build association prediction models, but most of them are through biological experiments, which are time-consuming and expensive. Therefore, it is necessary to develop a computational method focusing on microbe-drug spatial information to predict the microbe-drug associations. In this work, we use the biological information to construct heterogeneous networks of drugs and microbes. We propose a new method based on Multiple Kernel fusion on Graph Attention Network (GAT) to predict human microbe-drug associations, called GATMDA. Our method extracts multi-layer features based on GAT which can learn the embedding of microbes and drugs on each layer and achieve the purpose of extracting multiple information. We further fuse multiple kernel matrices based on average weighting method. Finally, combined kernel in the microbe space and drug space are used to infer microbe-drug associations. Compared with eight state-of-the-art methods, our method receives the highest AUC and AUPR on MDAD and aBiofilm dataset. Case studies for Human immunodeficiency virus 1 (HIV-1) and adenovirus further confirm the effectiveness of GATMDA in identifying potential microbe-drug associations.

## 1. Introduction

The human microbe is a complex and diverse community that has a very important impact on human health. Studies have shown that there are about bacterial cells in an adult’s body, equivalent to 10 times the size of human cells, and these cells can produce a large number of gene products to support various biochemical or metabolic activities in our body [1]. Microbes play an important role in human health, and changes in the human environment, such as diet, season, smoking, and vitamin injection, etc. [2-4]. It may lead to changes in the transcriptome, proteome and metabolome, which can further damage human tissues and eventually cause various diseases, such as obesity, diabetes, inflammatory bowel disease and cancer. Several studies [5-7] have shown that microorganisms participate in drug absorption and metabolism, thereby regulating drug efficacy and drug toxicity. Therefore, microbe-drug association research has attracted more and more attention.

A large number of microbe-drug potential relationships have been verified by traditional experiments. For example, Kovac et al. [6] demonstrated that the Enterococcus faecalis and Candida albicans strains were slightly susceptible to Ciprofloxacin. Szczuka et al. [7] showed that Ciprofloxacin inhibits the formation of Staphylococcus epidermidis biofilms. However, conventional wet-lab experiments to reveal microbe-drug associations are time consuming, laborious and expensive.

Thus, computational methods that effectively and accurately predict microbe-drug associations are useful additions to limited experimental approaches [14-17]. In 2017, Sharma et al. [14] developed a computational method to predict metabolic enzymes and gut bacterial species that can perform drug molecular bio-transformation. In the same year, Zhu et al. [16] proposed a microbe-drug prediction model HMDAKATZ based on KATZ measurement. Long [17] proposed GCNMDA, a microbe-drug association prediction framework based on GCN, which applied conditional random fields to ensure that similar nodes have similar representations. Although many calculation methods have been proposed to predict microbe-drug associations, the comprehensive features of microbes and drugs cannot be preserved during feature extraction.

To address the above issue, we propose a novel graph attention network, named GATMDA for microbe-drug associations prediction in a bipartite network. First, we quantify comprehensive features for microbes and drugs by assembling drug chemical information, microbe gene information and Gaussian interaction profile features. Secondly, we introduce graph attention networks with talking-heads, a variant of benchmark GAT, to learn node representations, which enables the model to preserve more informative representations. Third, we perform multi-core fusion of multiple kernels and matrices extracted by GAT in two spaces, respectively. Finally, a decoder is fed by the learned latent representation and outputs the probability standing for the presence of microbe-drug associations. Experimental results on three data sets, MDAD, aBiofilm and Drug Virus, indicated that our proposed model of GATMDA consistently outperformed eight state-of-the-art methods. Case studies on two diseases, HIV-1 and adenovirus, further validated the effectiveness of GATMDA.

The proposed GATMDA method has the following advantages: (i) The GATMDA method focuses on exploring the microbe-drug space information, by combining GAT and multiple kernel fusion. And by constructing a Heterogeneity network between microbes and drugs, the GATMDA effectively integrates the rich biological information between microbes and drugs. (ii) The GATMDA algorithm balances the regularization term by introducing a decay factor to dual Laplace regularized Least Squares, which further improves the prediction performance. (iii) Based on the GATMDA, we apply the method to three commonly used datasets, the prediction results reveal that the proposed approach outperforms existing methods in terms of AUC, and AUCR. (iv) Case study on HIV-1, only two of the top 30 drugs have no literature support, which further demonstrates the effectiveness of GATMDA in detecting microbe-drug associations.

## 2. Materials and Methods

### 2.1 Data sources

In order to speculate the new association in the human microbe-drug network, we can see it as a biological binary network prediction. We can represent the microbes and the drugs as two different types of nodes in the network. Define the node set of drugs N_d_ as 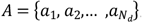. The node set of microbes *N*_*m*_ is described as 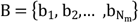. The edges in the network are the associations between microbes and drugs, which can be expressed as adjacency matrix 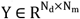. When Y_i,j_ = 1, a microbe b_j_(1 ≤ j ≤ N_m_) corresponds to a drug a_i_(1 ≤ i ≤ N_d_). Conversely, *Y*_*i,j*_ = 0 means that the association is unknown. Our goal is to obtain a prediction matrix of the same size as Y to predict unknown associations F^*^.In this study, we designed a multi-core fusion model (GATMDA) based on the graph attention network to infer new microbe-drug associations. Fig 1 shows the flow diagram of the GATMDA algorithm.

**Fig 1.**
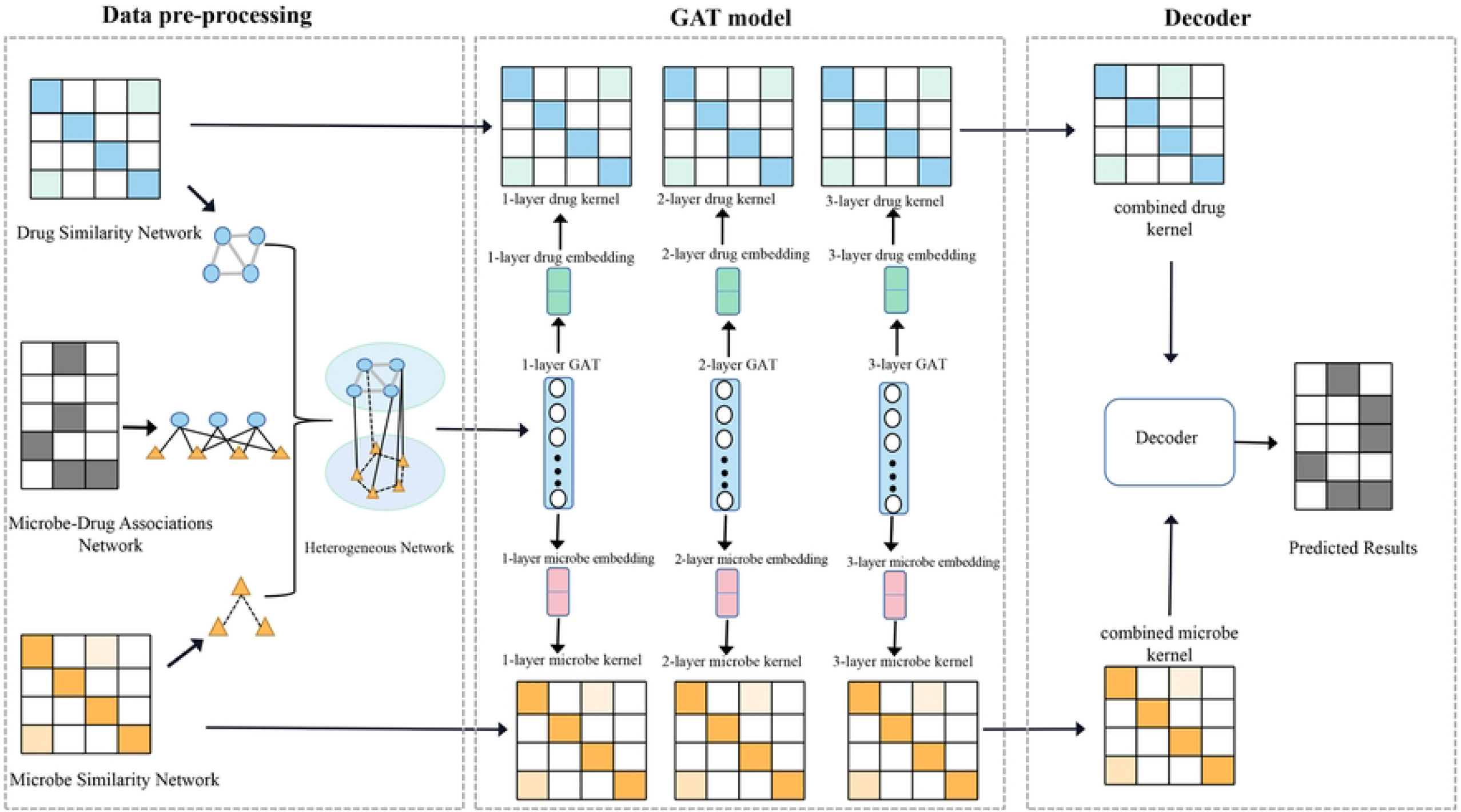
The schematic illustration of the GATMDA algorithm. GATMDA consists mainly of three modules. The first module is to construct input signatures for microorganisms and drugs datasets. The second module is designed to learn node representations based on GAT with speaking heads. The third module aims to perform multi-kernels fusion of multiple kernel matrices extracted from GAT and use a decoder to predict microbe-drug associations.

We use three datasets of known microbe-drug associations, the first is the MDAD dataset, which after removing redundant information. It includes 2470 known associations for 1373 drugs and 173 microbes. The second dataset is the aBiofilm, stores the resources of antibiofilm agents and their potential impact on antibiotic resistance. The aBiofilm has 23 unique types of anti-biofilm preparations, including 1720 unique anti-biofilm preparations (drugs) and 140 unique targeted microorganisms. We ultimately select 2884 microbe-drug associations for our study. The last is the DrugVirus dataset, which records the activity and development of compounds related to multiple human viruses, including Adenovirus. In this dataset, we screen 933 relevant associations from 95 viruses and 175 drugs for our experimental data.

Table 1 shows the detailed data volumes for the above three datasets. In this paper, we use these three datasets to test the predictive power of our model.

**Table 1.** Detailed data for three datasets.

### 2.2. Construction of the Heterogeneous network

To incorporate network information into data integration, we construct a heterogeneous network that includes microbial network S_m_, drug network S_d_, and association network between microbes and drugs. The details about the calculations of S_m_ and S_d_ can be found in Supplementary Materials. Finally, a heterogeneous network Y defined by an adjacency matrix [18] is constructed:

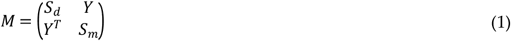

### 2.3. Graph attention neural network

The GAT is a space-based graph convolutional network [19]. The core of the graph attention mechanism is to focus on the feature contributions of more important neighbors in the process of aggregating neighbor features [20]. GAT is used in our research to extract microbe and drugs signatures. Specifically, for the adjacency matrix of the binary network defined above, GAT is defined as follows:

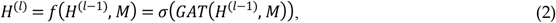

where H^(l)^ is the l-layer embedding of nodes, l = 1,…,L, σ(.) is a non-linear activation function. GAT represents a single graph attention layer, and the GAT architecture of the whole L layer is stacked by multiple graph attention layers. The initial input is a set of node features 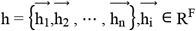, where n is the number of nodes and F is the number of features in each node. This layer will generate a new set of node features 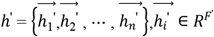 and we transform the input features into higher-level features using a learnable linear transformation by applying a weight matrix 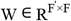 to each node. Then, we calculate the attention factor as:

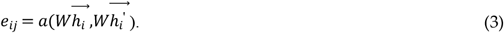

After normalization by the softmax function, we accept the coefficients as:

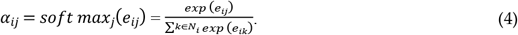

By substituting (3) into (4), the coefficients of attention mechanisms are as follows:

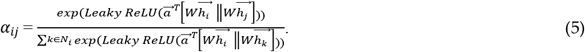

where, α is the attention coefficient, 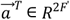 represents the parameterized weight vector, LeakyReLU is the activation function, T is the matrix transpose, || is the connection operation, and N_i_ is the set of neighbors of node i. After calculating the normalized attention coefficient, the final output features of each node can be calculated as:

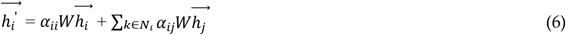

In our study, for the first layer, we constructed the initial embedding *H*^(0)^, as follows:

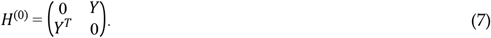

### 2.4. Multiple kernel fusion

The multi-layer GAT model can calculate multiple embeddings representing information with different graph structures. Since different embeddings represent different structural information, the kernels composed of different embeddings will represent the similarity between nodes of different angles. Combining the existing similarity matrix, we can obtain the kernel sets of drug space 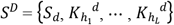,and microbe space 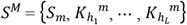. 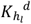 and 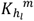 are the kernel matrix of the drug and microbial embedding in each layer, respectively. The details about the calculation of kernel matrix can be found in Supplementary Materials.

In order to improve the performance of predicting microbe-drug associations, we performed multi-kernel fusion of the above kernel in two spaces, respectively, which can combine multiple kernel matrices by weighted method. The combined kernel is defined as follows:

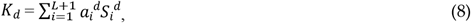

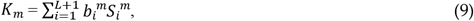

where *S*_*i*_^*d*^ and *S*_*i*_^*m*^ are the i-th kernel in the drug and microbial nucleus sets, respectively. *a*_*i*_ and *b*_*i*_ are the attention factors corresponding to each nucleus, and *L* is the number of layers.

### 2.5. Decoder for microbe-drug associations reconstruction

We apply an improved dual Laplace regularized Least Squares (DLapRLS) framework to predict associations, improving prediction performance. The loss functions are defined as follows:

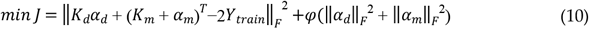

where, ‖.‖_F_ is the Frobenius norm, 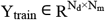 is the adjacency matrix of microbe-drug association in the training set; *α*_*d*_, 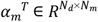 is the trainable matrix; 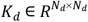 and 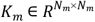 is fusion cores in two feature spaces, respectively. And *φ* is decay factor to balance the regularization term.

Therefore, the predicted combination of microbe-drug association from the two feature spaces is as follows:

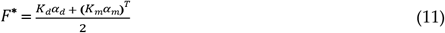

## 3. Results

We first introduce the selection of model parameters using 5-fold cross-validation (5-CV) on the MDAD dataset. Under the same conditions, the impact of different models can be tested for predictive analysis. Next, we use three cross-validation methods (2-CV, 5-CV, and 10-CV) on three datasets to compare with other existing methods.

### 3.1. Parameter sensitivity analysis

k-fold cross-validation (k-CV) is widely used to evaluate prediction performance. During cross-validation, all associations are divided equally into k parts. In each fold, one of them is selected as the test set, and the rest is used as the training set for training and validation of the model, for a total of k folds. In this method, we can calculate true positive (TP), false positive (FP), true negative (TN), and false negative (FN), respectively. In addition, we mainly use the following evaluation measures: area under the receiver operating characteristic curve (AUC) and area under the precision recall curve (AUPR). In our method, there are several important parameters, such as the decay factor φ, the iteration time N, learning rate and the kernels in graph embedding. In this section, all experiments are conducted based on dataset MDAD and are evaluated under 5-fold CV. In addition, we set the number of layers L = 3, and the embedding dimension K_1_ = 256, K_2_ = 64, and K_3_ = 32.

In this study, we use the delay factor *φ* to control the contribution of regularization terms in (10). In our experiment, we vary the *φ* from 0.000005 to 0.5 with a step value of 10. From Fig 2 (A), we can conclude that this parameter has a relatively slight effect on performance, which indicates that our model is robust to the delay factor *φ*. The best performance is achieved when φ = 0.0005. The learning rate is also a very important parameter. Choosing an appropriate learning rate is challenging, because the model is difficult to converge when the learning rate is too large, and a small value may result in a long training process. A reasonable learning rate can allow the model to converge to the local minimum. Therefore, we vary the learning rate from {1e−1, 1e−2, 1e−3, 5e−3, 1e−4, 5e−4}, and we evaluate the performance of GATMDA at these different learning rates. The results are shown in Fig 2B. We can observe that from 1e−1 to 5e−4, the performance of GATMDA first increase and then decrease slightly. GATMDA works best when the learning rate is 1e−3.

**Fig 2.**
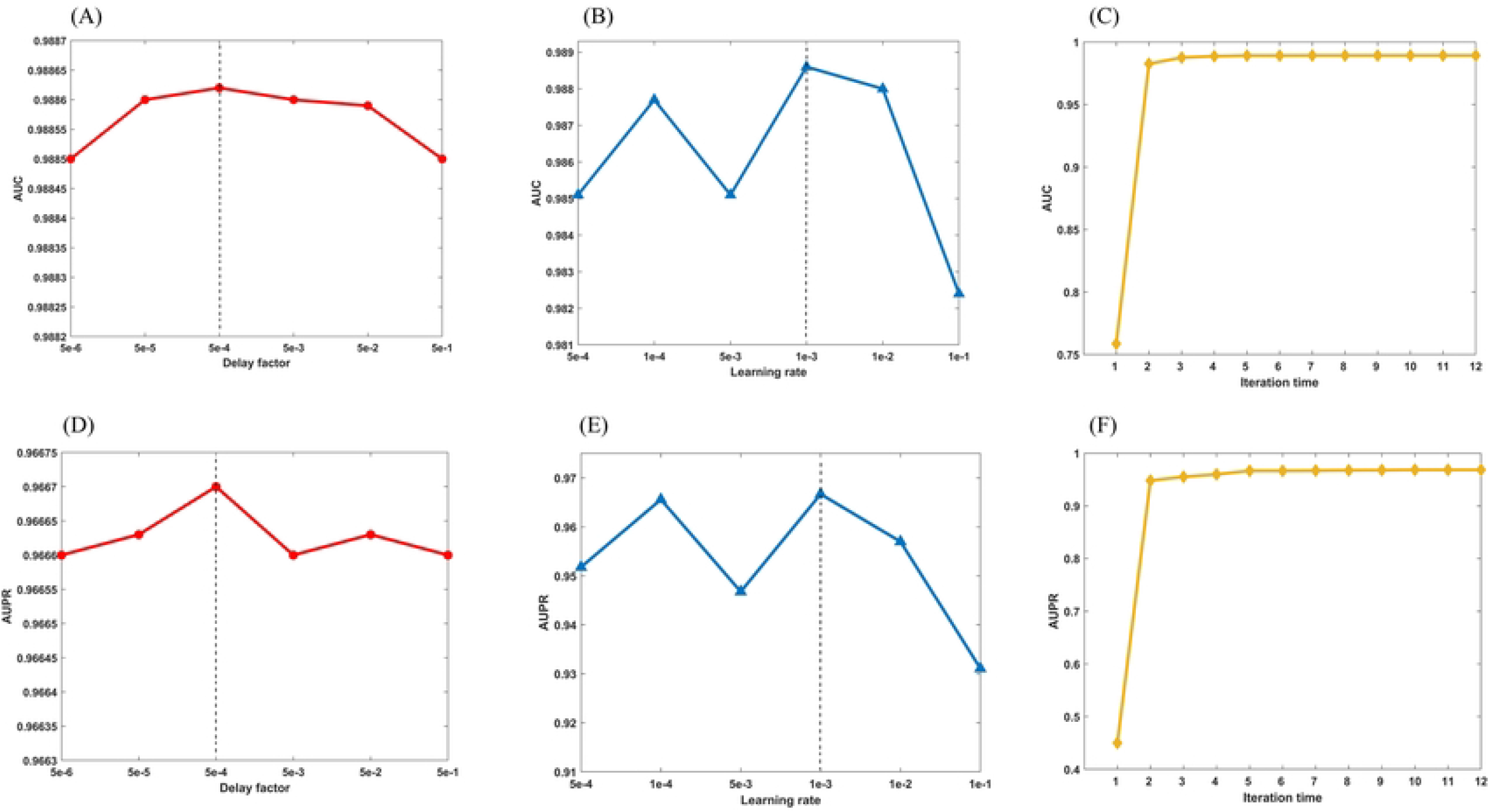
Parameter sensitivity of GATMDA

In addition, the number of iteration number N plays an important role in our model. The iteration number N controls the number of updates to trainable parameters. We range N from 1 to 12 with a step value of 1. Fig 2F shows the AUPR under different iterations. As shown in Fig 2F, it can be observed that the AUPR value tends to stabilize when the number of iterations is 5. To make the model fully convergent, we choose the iteration number of 10.

In our model, GAT is used to extract microbe and drug signatures. The multi-layer GAT computes the embedding of different layers and fuses multiple kernel matrices based on graph embedding information and initial similarity. In this section, we discussed initial similarity measures, kernel matrices generated by different layers, and the impact of combined kernels on association prediction. Since the three layers GAT were applied in our study, we used h_l_ +GAT MDA(l = 1,2,3)to represent the kernel matrix obtained by the GATMDA model in each of the three layers. GATMDA is a model that assigns average weight to the above three kernel matrices. The results are shown in Fig 3. As can be seen from the Fig 3, the AUC value of h2+GATMDA is better than that of h1+GATMDA and h3+GATMDA, and the AUPR value of h1+GATMDA is better than that of h2+GATMDA and h3+GATMDA, which means that each kernel matrix generated by GAT provides useful information for prediction. On the other hand, comparing GATMDA and single-kernel model, GATMDA performs significantly better than single-kernel model, which indicating that multi-kernel model can combine more information to make predictions. Therefore, in our study, using attention mechanisms for information fusion can improve predictive performance.

**Fig 3.**
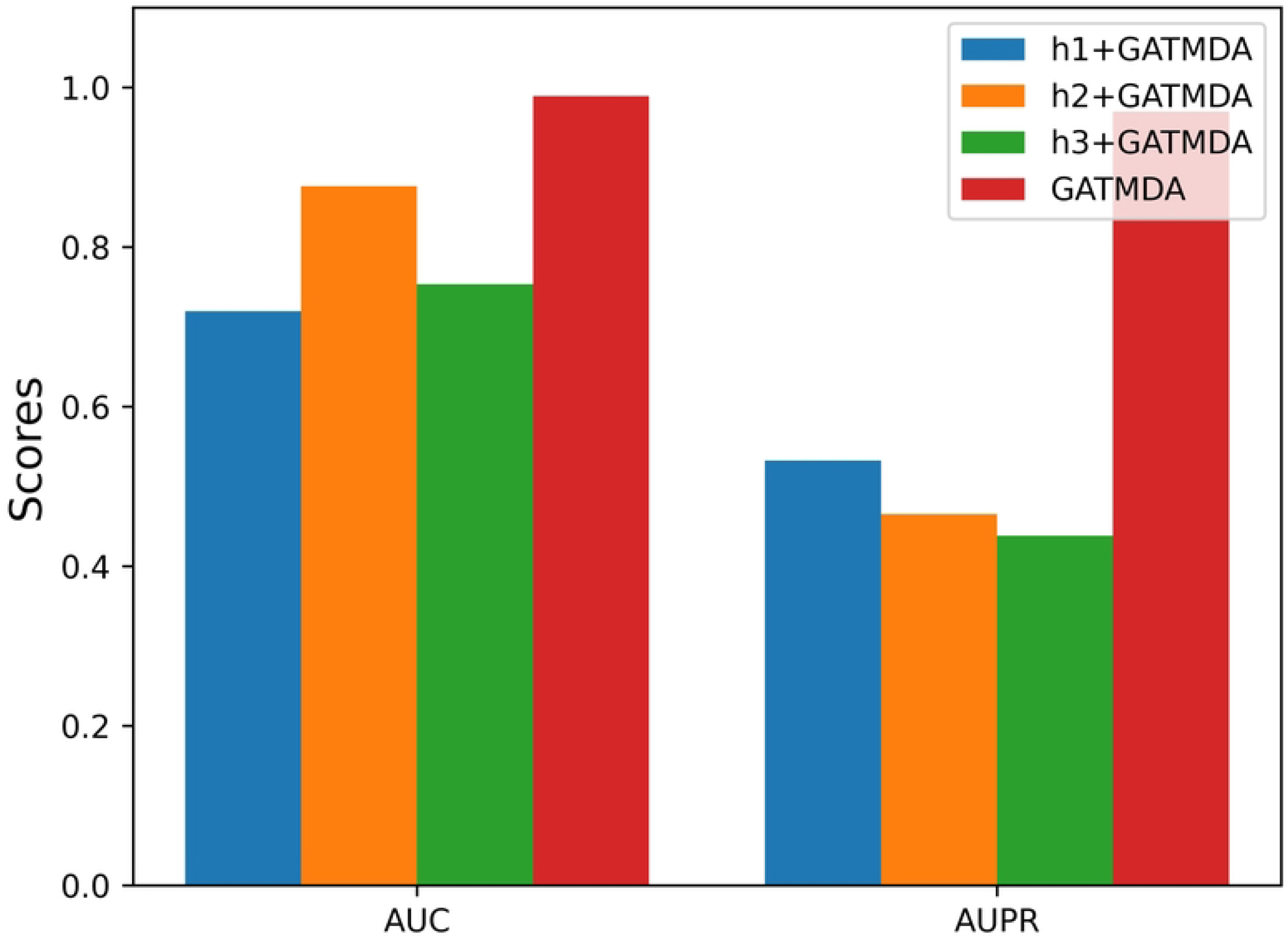
Performance of GATMDA based on different kernels.

### 3.2. Comparison with existing excellent tools

We compare this method with existing biological binary network prediction methods. For example, KATZHMDA, WMGHMDA, and NTSHMDA are used to predict the correlation of microbial diseases. IMCMDA and GCMDR are used for microRNA-disease association prediction and identification of miRNA-drug resistance relationships, respectively. BLM-NII is developed to address drug-target interactions. MKGCN and SCSMDA are used to predict the associations of microbe-drug. Among them, SCSMDA [21] is the latest method to predict microbe-drug association based on structure-enhanced contrast learning and self-paced negative sampling strategy.

In this paper, we present a GATMDA model to predict microbe-drug associations. Our model learns advanced feature representations from raw input graphs and node attributes via GAT. We plot the potential representation of all drugs into a two-dimensional space in the MDAD dataset using the t-SNE tool. The visualization results are shown in Fig 4. As shown in Fig 4A, the scatter plot shows the distribution of drugs in the original space. It can be seen that the drug distribution is in a confusion state before training the drug, while the drug shows a clear clustering pattern through the underlying representations learned by our model, as shown in Fig 4b.

**Fig 4.**
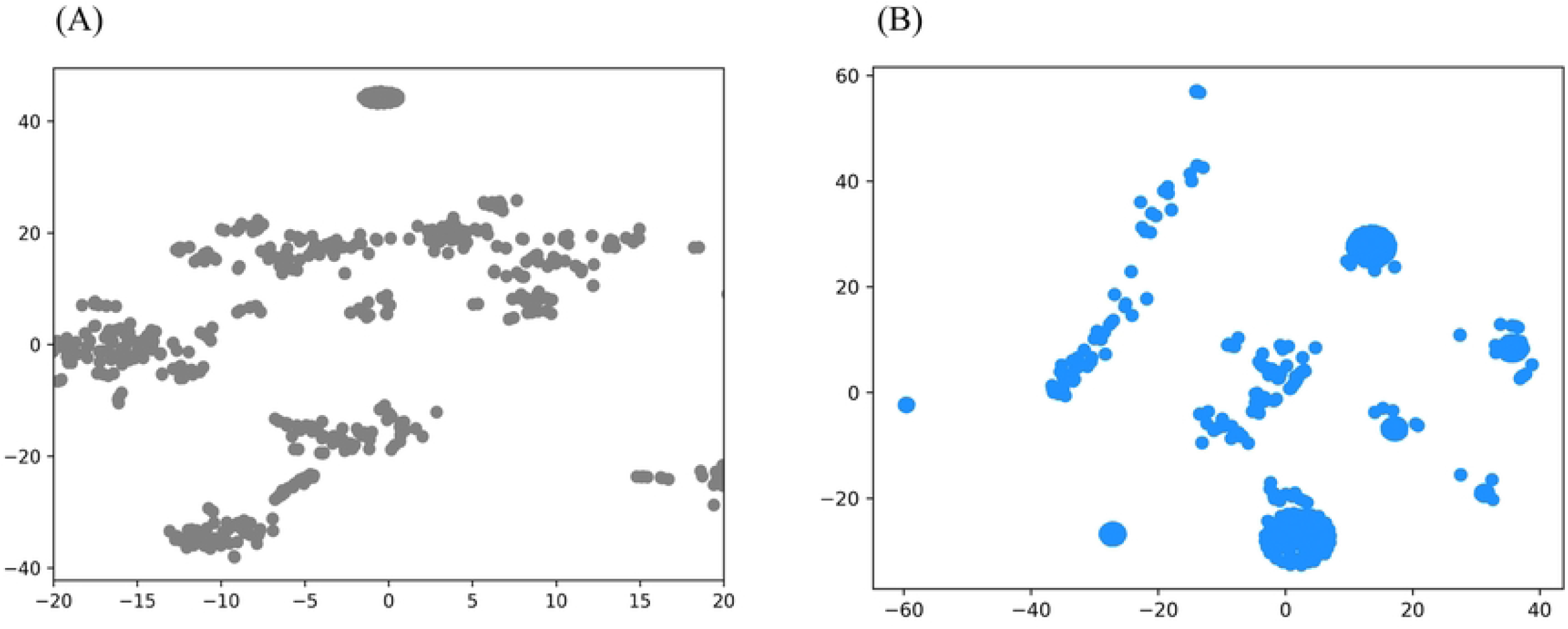
Potential representation of drugs in MDAD datasets learned by t-SNE tools. (A) Drug distribution visualized in the original attribute space. (B) Drug distribution is visualized in the learned latent representation space.

To make a fair comparison, we run these methods on the default parameters of the MDAD dataset and perform a 5-fold cross-validation for all methods. Table 2 and Fig 5 shows the results of 5-CV on MDAD dataset. Among all the methods, our proposed GATMDA model has the best predictive performance, with an average AUC of 0.9886 and an average AUPR of 0.9667. In terms of AUC and AUPR, it outperforms the remaining 8 models, respectively. To further evaluate the validity and robustness of our model, we also performed GATMDA and all baseline methods on two additional datasets, aBiofilm and DrugVirus. The GATMDA model achieves the best AUC and AUPR on the aBiofilm dataset (AUC: 0.9941, AUPR: 0.9869). However, the DrugVirus dataset (AUC: 0.9836, AUPR: 0.8795) have the best AUC and the second highest AUPR. This result may be due to less correlation with the dataset.

**Table 2.** Performance comparison between baseline methods and our method on datasets MDAD, aBiofifilm and DrugVirus under 5-fold CV.

**Fig 5.**
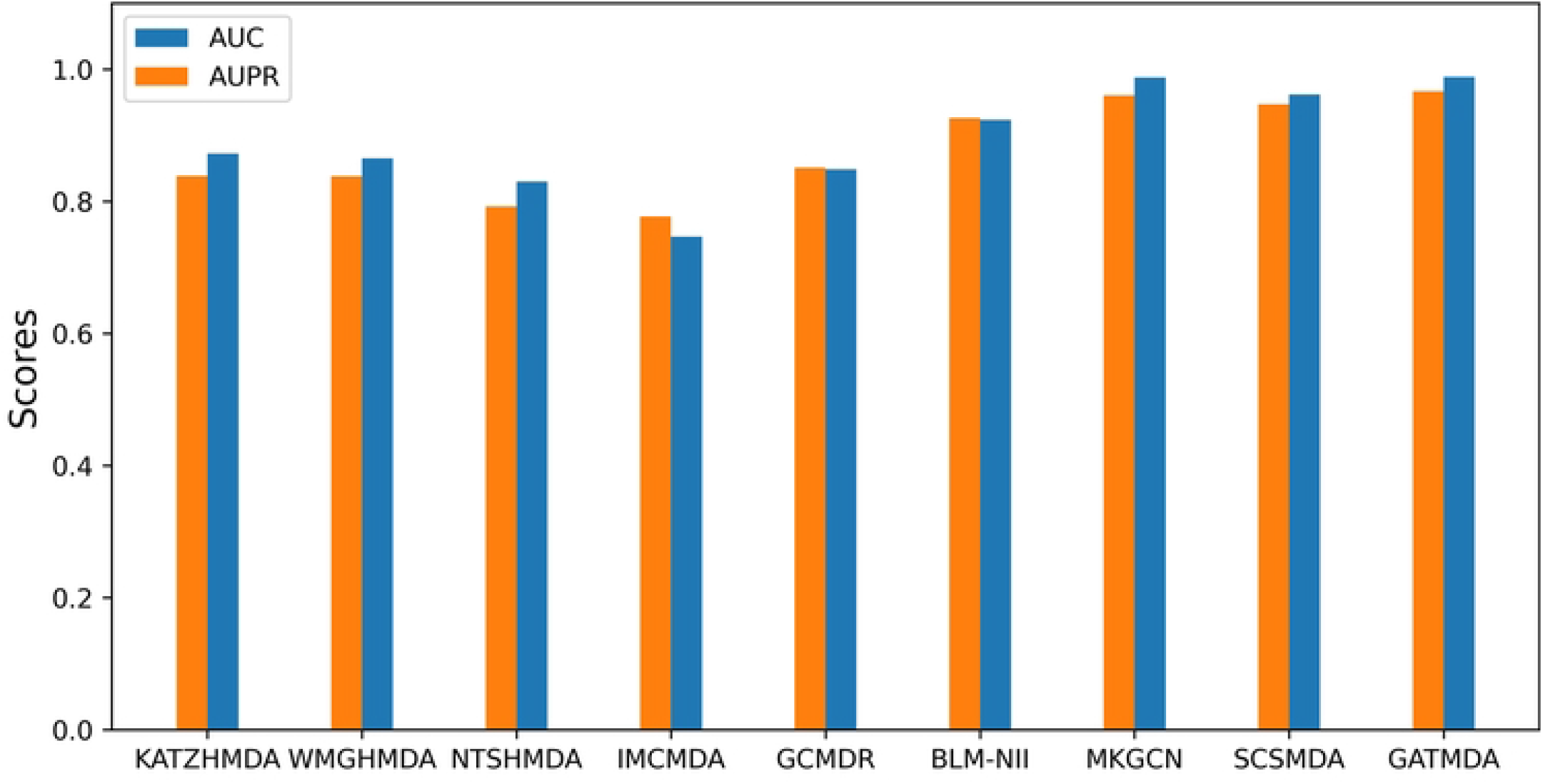
Performance comparisons of GATMDA with eight competitive methods on MDAD datasets.

In addition, we also perform comparative experiments on these three datasets for all methods under 2-CV and 10-CV settings, as shown in Supplementary Tables S1 and S2. The results confirm that GATMDA outperform all 8 state-of-the-art methods on the first two datasets, indicating that GATMDA is an efficient and robust computational model for predicting microbe-drug associations. We demonstrate the ROCs and PRs of prediction performance for the above three datasets with three cross-validation methods. The ROC and PR curves based on three cross-validations are illustrated in Fig 6.

**Fig 6.**
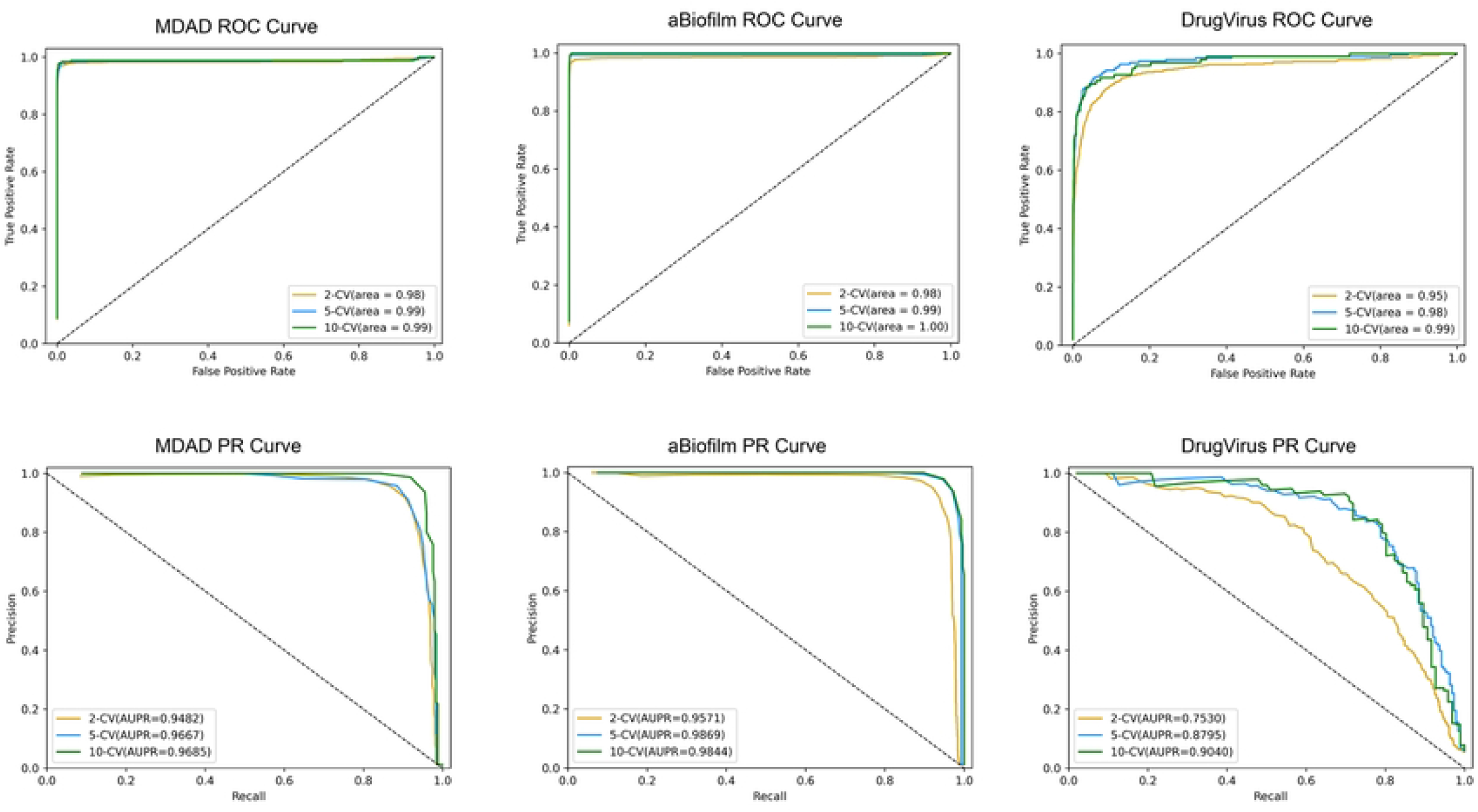
The ROC and PR curves of GATMDA on the MDAD, aBiofilm and DrugVirus datasets.

### 3.3. Case Studies

In this section, we further test the prediction effect of GATMDA through two case studies based on the DrugVirus dataset. We choose HIV-1 and Adenovirus as our case studies to test the predictive performance of our model while and predict drugs that might be effective therapeutically.

HIV is a retrovirus that destroys CD4 T cells and is the causative agent of acquired immunodeficiency syndrome (AIDS). HIV is divided into 2 types, both of which cause AIDS: HIV-1, which causes a global epidemic; HIV-2, which is weakly pathogenic, is mainly confined to West Africa. Thus, we select HIV-1 for the case study experiment. In the experiment, Griffith, B P et al. [22] measured the antiviral efficacy of stavudine (2’,3’-didehydro-3’-deoxythymine) against HIV-1 in 15 HIV-infected patients. The test results show that stavudine has a significant and long-lasting antiviral effect. Enfuvirtide (ENF), a novel HIV-1 fusion inhibitor, has potent antiviral activity against HIV-1 both in vitro and in vivo. And Enfovirtide is the first licensed inhibitor of HIV entry [23]. For HIV-1, we select the top 30 of the 175 predicted drugs to test the effectiveness of GATMDA (Fig 7a). Also, we plot a bar graph and dot plot to visualize the top 30 predicted drugs (Fig 7b-c). As shown in Table 3, the predicted HIV-1-related drugs, all of the top 10 drugs are supported in the literature. Among the predicted top 20 and 30 drugs, 95% and 93% of the drugs are supported by the literature and proved to be possible for treating or preventing the HIV-1. These prediction results demonstrate the ability of the GATMDA model to predict potential associations in the microbe-drug network.

**Table 3.** Top 30 predicted HIV-1-associated drug.

**Fig 7.**
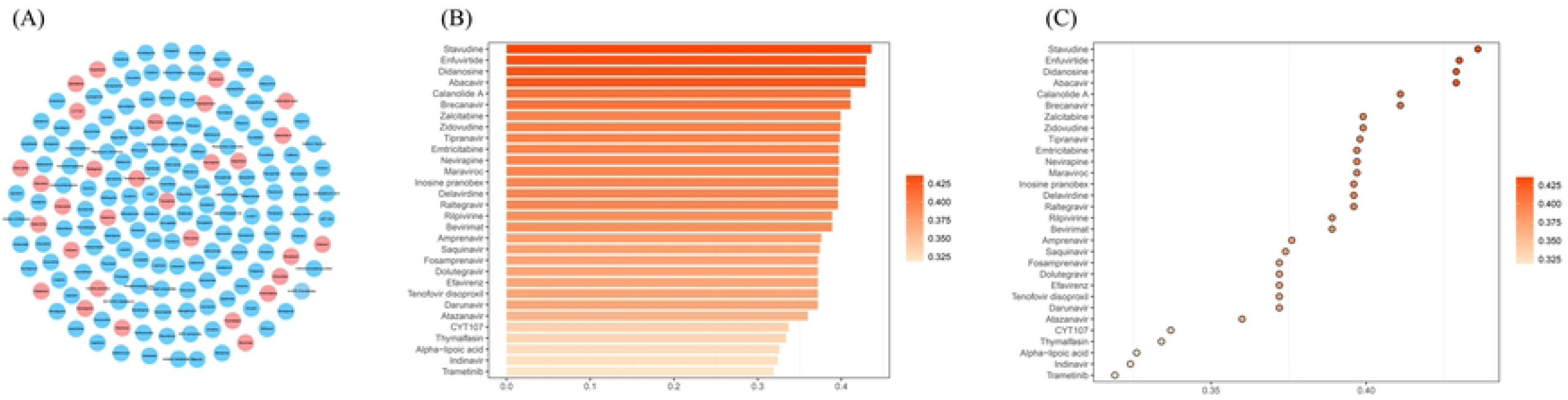
Visualization of the top 30 predicted HIV-1-related drugs. (A) Top 30 of the 175 drugs predicted by GATMDA. (B) Bar graph of the predicted top 30 drugs. (C) Dot plot of the predicted top 30 drugs.

Adenovirus was originally isolated from tissue culture of human adenoid proliferators. In humans, adenovirus causes acute mucous membrane infections of the upper respiratory tract, eyes, and regional lymph nodes, very similar to the common cold. Adenovirus can also cause epidemic keratoconjunctivitis (EKC). Supplementary Tables S3 shows the drug names and literatures. For example, the anti-AdV5 activity of CDV is mediated by viral DNA Pol, and their antiviral activity may occur by (non-obligate) strand termination and direct inhibition of AdV5 Pol activity (at high concentrations) [24]. Brincidofovir is an orally bioavailable lipid conjugate of cidofovir that has in vitro activity against adenoviruses [25]. Among the top 20 drugs predicted to target Adenovirus, 70% of drugs are supported by the literature.

## 4. Discussion and Conclusions

The microbes that inhabit and on the human body play a key role in human health. Predicting microbe-drug associations can facilitate the effective development of drugs and personalized medicines, and understand the interconnected links between microbes and drugs. Traditional culture-based methods are time-consuming and expensive. Compared to traditional methods, computational methods are able to identify target existing drugs or target microbes for new drugs with known microbes on a global scale.

In this paper, we propose a computational framework GATMDA to discover microbe-drug associations. Our framework GATMDA consists of two parts. In the first part, we use the GAT for feature extraction. Experiments show that more reliable inference information can be generated using this mechanism. The other part is the improved DLapRLS for prediction, which fully utilizes the information of microbe-drug space for prediction. Different from traditional multi-kernel learning, we construct kernel matrices by extracting various embedding features through multi-layer GAT, which can provide various kernel matrices and achieve the purpose of using multiple information. In the experiment, the GATMDA model has excellent performance on three existing microbial-drug association datasets. In addition, we conduct a case study of HIV-1. This case study shows that GATMDA can accurately discover new microbe-drug associations.

Although the GATMDA algorithm proposed in this paper has an excellent prediction performance, there is a certain deviation for datasets with different densities. For example, GATMDA performs weaker on the DrugVirus dataset than on the MDAD and aBiofilm datasets, indicating that there is still room for improvement in the generalization performance of GATMDA models. In addition, as mentioned earlier, the influence of microbes on the course of drug treatment includes activation, passivation, and toxicity. Accurately identifying the type of unknown microorganisms on drugs is a basic requirement in drug development and precision medicine, but the GATMDA algorithm cannot predict the type of microbe-drug. Therefore, in order to more accurately understand the mechanism of microorganisms in the drug treatment process, the development of an effective deep learning model to predict the relationship between microbe-drugs requires further research.

## Supplementary Materials

The following supporting information can be downloaded at: www.mdpi.com/xxx/s1.

## Author Contributions

Conceptualization, S.S. and S.K.; methodology, S.S.; software, S.S., S.K.; validation, S.S. and Q.Z.; formal analysis, Q.Z.; investigation, S.K.; resources, J.Z.; data curation, S.K.; writing—original draft preparation, S.S. and S.K.; writing—review and editing, J.Z.; visualization, S.S. and S.K.; supervision, J.Z.; project administration, J.Z., S.S.; funding acquisition, J.Z. All authors have read and agreed to the published version of the manuscript.

## Institutional Review Board Statement

Not applicable.

## Informed Consent Statement

Not applicable.

## Data Availability Statement

The datasets and corresponding codes are available at https://github.com/SSRHallow/GATMDA.

